# A drop-on-demand bioprinting approach to spatially arrange multiple cell types and monitor their cell-cell interactions towards vascularization based on endothelial cells and mesenchymal stem cells

**DOI:** 10.1101/2022.07.20.500797

**Authors:** Joshua Weygant, Fritz Koch, Katrin Adam, Kevin Troendle, Roland Zengerle, Günter Finkenzeller, Sabrina Kartmann, Peter Koltay, Stefan Zimmermann

## Abstract

Spheroids, organoids, or highly-dense cell-laden droplets are often used as building blocks for bioprinting, but so far little is known about the spatio-temporal cellular interactions post printing. We present a drop-on-demand approach to study the biological interactions of such building blocks in micrometer dimensions. Droplets (containing approximately 700 cells in 10 nl) of multiple cell types are patterned in a 3D hydrogel matrix with a precision of less than 70 μm. It is applied to investigate interactions of cell types relevant for vascularization approaches. We show that a gap of 200 μm between droplets containing endothelial cells (HUVECs) and adipose-derived mesenchymal stem cells (ASCs) leads to decreased sprouting of HUVECs towards ASCs and increased growth from ASCs towards HUVECs. For mixed aggregates containing both cell types, cellular interconnections of ASCs with up to approximately 0.8 millimeter length and inhibition of HUVEC sprouting are observed. When ASCs are differentiated into smooth muscle cells (SMCs), HUVECs display decreased sprouting towards SMCs in separate aggregates, whereas no cellular interconnections or inhibition of HUVEC sprouting are detected for mixed aggregates. These findings demonstrate that this approach acts as a new tool to investigate cell-cell interactions of different cell types in 3D bioprinted constructs.

## 1. Introduction

Vascularization plays an important role in tissue engineering applications aiming at the development of artificial tissues substitutes. In general, a microvascular network is essential to maintain healthy tissue, prevents cell death and allows the tissue to function correctly due to waste product removal and supply with nutrients and oxygen. ^[1]^ Therefore, a vascular network is required for bioengineered constructs to ensure correct functioning of the tissue, to prevent cell death in constructs that are thicker than 200 μm, and to enable long-term viability. ^[2]^ Several bioprinting approaches aim to create a vascular network, for example via sacrificial printing, coaxial deposition or self-assembly of cells post printing. ^[2,3]^ In this context, endothelial cells (ECs), such as human umbilical vein endothelial cells (HUVECs) and smooth muscle cells (SMCs) are often used together to fabricate blood vessel like structures. ^[4–7]^ Additionally, adipose-derived mesenchymal stem cells (ASCs/MSCs) are also used, because they are known to be able to secrete vascular endothelial growth factor (VEGF), an angiogenic growth factor, and it has been demonstrated that they can be differentiated into ECs and SMCs *in vitro*. ^[8–10]^ Since MSCs can be differentiated into several cell types, they are attractive candidates for several tissues. For example, they can be differentiated into osteogenic cell types, making them attractive candidates for bone tissue engineering. ^[11]^ Additionally, HUVECs have been bioprinted with ASCs and the printed scaffold led to blood vessel formation as well as well as synthesis of a calcified bone matrix *in vivo*. ^[12]^ A lot of research has been devoted to better understand the interactions between these cell types, but they are still not fully understood. For example, it has been demonstrated that ASCs support angiogenesis and promote lumen formation, whereas it has been also shown that co-culture of ASCs and HUVECs does not enhance angiogenic potential of HUVECs. ^[13,14]^

In 3D, spheroids are often used to investigate cell behavior since they mimic the natural cell environment more accurately than cells in 2D environments. ^[15]^ For spheroids in hydrogels, active sprouting, migration and enhanced differentiation potential have been demonstrated. ^[16–18]^ Within spheroids, the cells interact with cells from neighboring spheroids through the secretion of growth factors or through physical contact, for example between sprouting/migrating cells. ^[19]^ Thus, the distance between spheroids is a crucial parameter for artificial tissues. However, there are few techniques that allow to control the spatial arrangement and the distance between two or more spheroids in 3D environments. Controlling the distance between the spheroids allows to investigate cell-cell interaction between spheroids of different cell types better and few approaches have been developed. There is the Kenzan method, which uses microneedles called “kenzans” to align spheroids, placement of spheroids based on aspiration, and recently micropatterning of hydrogels based on a stamp technique and subsequent placement of spheroids into resulting gaps have been reported.^[18,20,21]^

Previously, we have developed a printing process to precisely position HUVECs in a fibrin hydrogel and have shown that this approach can be used to compare highly-dense cell-laden droplets of HUVECs with HUVEC spheroids. ^[3]^ Also, in this study it was revealed, that a droplet-based approach yields longer cumulative sprout lengths (CSL) than spheroids, indicating a higher level of interaction between the cells. So, the use of such droplets of high-density cell suspensions could exhibit similar or even enhanced functional properties of spheroids like the sprouting behavior indicated above for endothelial cells, but also the self-assembly capacity of renal epithelial cells for the formation of tubular structures. ^[22]^ Therefore, a high-density cell droplet approach could offer an alternative to the laborious spheroid manufacturing process and would enable the direct use of the cells after harvesting in a single-step biofabrication process.

Based on this work, for the present study we developed a new method that allows to print highly-dense cell-laden droplets of multiple cell types in a 3D environment with high precision towards each other. Droplets containing different cell types are printed onto a fibrin hydrogel and are embedded into a sandwich-like structure to encapsulate the printed aggregates in a 3D environment. With the developed method it is possible to print different cell types in various constellations, from a full overlap up to increasing distances. The patterns are simply changed by adjusting the G-Code. First, the method was established with immortalized mesenchymal stem cells (iMSCs) and was then expanded by additional cell types like HUVECs, ASCs, and ASCs differentiated towards SMCs, to investigate the influence of the cell types on each other. We printed HUVECs and ASCs via individual or mixed cell suspensions in various constellations. Afterwards, ASCs were substituted with SMC-differentiated ASCs (dASCs) and the experimental setups were repeated to investigate the influence of the aggregates onto each other and to compare the behavior between HUVECs and ASCs

## 2. Results and Discussion

### 2.1. Process to print different cell types with high precision on the same substrate

The developed process is described briefly and can be found in more detail in the method and materials section (compare 3.1.). It is schematically shown in **Figure 1**. The first cell type is printed onto the fibrin substrate with a pattern that is pre-defined via G-Code. Afterwards, a second cell type is printed onto the fibrin substrate containing the first cell type and printing patterns are designed in relation to the first cell type. Then, a second layer of fibrin is added on top, and cells are incubated in cell culture medium for several days. The samples can be then analyzed, for example via antibody staining or via visual analysis.

**Figure 1:**
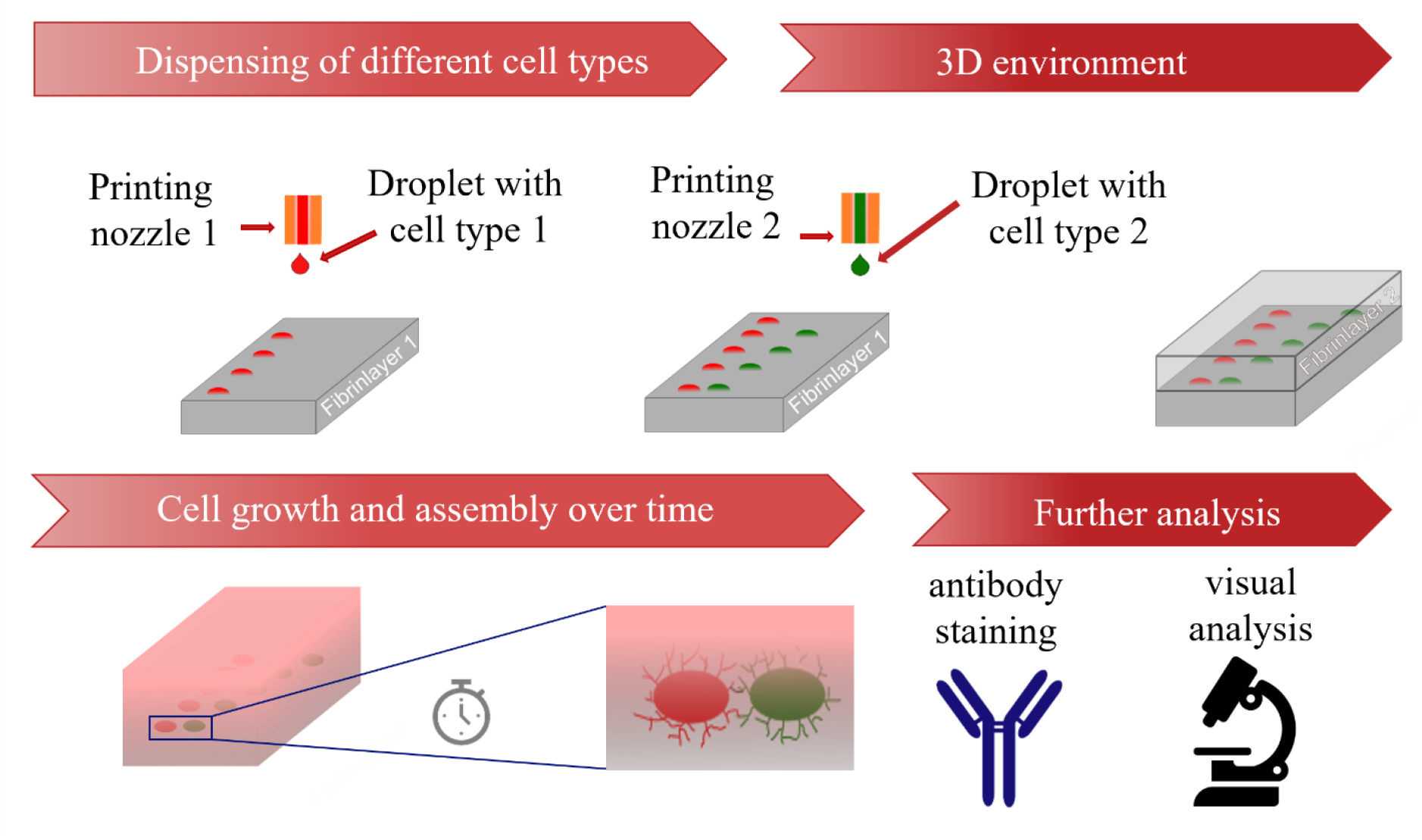
Schematic representation of the developed process from printing process to analysis.

Within one experiment between N = 3 - 15 samples with n = 6 - 48 cell aggregates were fabricated via the described process, demonstrating the capability to pattern numerous hydrogels with high number of cell-aggregates. A droplet-based technique displayed the advantage of printing complex patterns of cell aggregates with defined patterns and little effort in preparation. Compared to approaches with spheroids, the cells were directly harvested from the culturing flasks, suspended into the bioink and printed. Also, in contrast to other methods, the printed patterns can be changed easily by adjusting the G-Code. This allows flexible adjustments within one experiment. For this work, the cell types used are relevant for engineering vascularized tissue. However, the established method can be applied to various cell types and to more than two cell types.

### 2.2 Printing Performance

Several experiments have been performed to assess the printing performance in terms of robustness, precision, and cytocompatibility (impact on cell viability). First, patterns containing arrays of 6 × 8 iMSC cell aggregates with pitches (distance from center to center from the cell aggregates) of 500 μm between them were fabricated on N = 3 samples (**Figure S1**). The aggregates had a circular shape and a radius of r = 125 ± 5 μm before covering. After adding the second fibrin layer, a radius of r = 126 ± 6 μm was measured. Also, an overlay of images from before and after covering showed no displacement or disintegration of the cell spots. Thus, it was concluded that adding a second layer does not influence shape fidelity of cell aggregates.

Second, a live/dead assay was performed with iMSCs post-printing to characterize the influence of the dispensing technique onto cell viability by varying the stroke of the piezo-actuator or the stroke velocity of the piezo-actuator. For a constant stroke of 21.6 µm the stroke velocity was varied from 30 to 120 μm/ms in 10 μm/ms steps and for a constant stroke velocity of 60 μm/ms, the stroke was increased from 10.8 µm to 36 µm in 3.6 µm steps. For each measurement point n = 10 cell spots were analyzed. The parameters to obtain droplets with a volume of 10 nl yielded viability rates around 80%, which was in range of the positive control (approx. 85 % viability). Results are shown in **Figure S2**. The influence of an analogous drop-on-demand bioprinting approach onto viability of HUVECs was already investigated previously. ^[3]^

To determine the precision of the developed process, iMSCs were stained with Celltracker™ (ThermoFischer Scientific, C34552, Waltham, MA, USA) green and red, which was representative for two different cell types in future experiments. A pattern (**Figure 2 a**) was printed, where red iMSCs were printed in a line with a pitch of p_x,r_ = 1000 μm and p_r,y_ = 0 μm between the aggregates. Afterwards, green iMSCs were printed in relative position to the red iMSCs, where the green iMSCs had an ideal pitch of p_x,g_ = 0 μm and p_y,g_ = 800 μm from the respective red iMSC cell aggregate (indicated by the yellow circles in **Figure 2 a**). N = 8 technical replicates with n = 21 - 32 cell pair aggregates were fabricated. Afterwards, the deviations Δx and Δy of each cell aggregate from their ideal position was analyzed. Results for the deviations of the red iMSCs from their ideal position are shown in **Figure 2 b**, where the symbol represents individual spots within a run and colors represent different runs. The average deviations in x- and y-direction were obtained to be 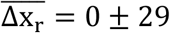 and 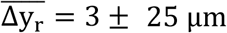.

**Figure 2:**
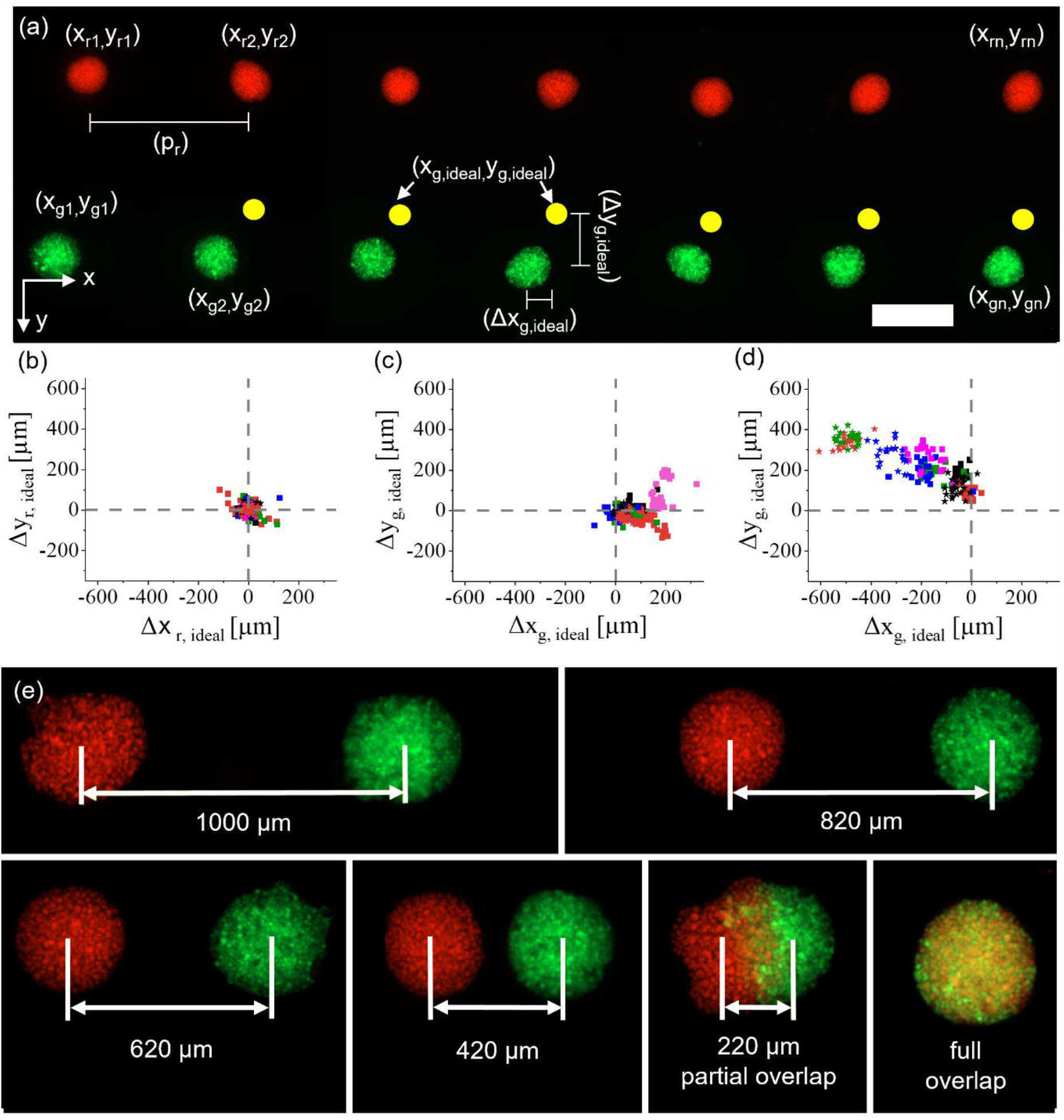
Characterization of the DoD bioprinting process developed for multiple cell types. Red- and green-labelled iMSCs were printed with a pre-defined pattern (a) and the deviation from the ideal position (yellow spots) was measured. Scalebar: 500 *µm*. The deviations of red iMSCs were analyzed (b) and then the deviations of green iMSCs when samples were not moved (c) and when samples were removed and retransferred to the printing platform (d) were analyzed as well. Dotted line shows the ideal position. (e) shows iMSCs printed with decreasing pitches in approximately 200 *µm* steps up to an entire overlap.

The location of the green cell spots was analyzed in two separate experiments. First, the glass slides with the hydrogels were not removed from the printing platform between printing the red and green iMSCs. This yielded average deviations from the ideal position of 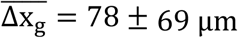 and 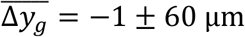. The results are shown in **Figure 2 c**. In the second experiment, the hydrogels were removed from the printing platform after printing the red iMSCs, transferred into a humid atmosphere and re-transferred onto the printing platform when the green iMSCs were printed. This process increased the deviations from the ideal position to 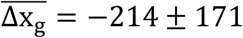 and 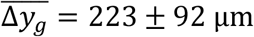. Results are shown in **Figure 2d**. The increase in average deviation is most likely caused by the transferring process of the samples. However, within a run, average alignment errors were ± 58 μm in x- and ± 54 μm in y-direction on the individual samples. This is also indicated by the clustering in the results in figure 2 c and d. Besides initial alignment errors, by monitoring the printing process after each print, it was possible to print defined intervals between the cell types, for example from a full overlap up to pitch of 1000 μm (**Figure 2 e**).

Based on these results, a printing process without transferring the samples between printing different cell types would yield lower errors. However, in order to prevent evaporation of the fibrin hydrogel, samples were temporarily stored in the humidified incubator between printing the two cell types. This issue could be addressed in further developments, e.g., by implementing an evaporation protection in the bioprinting device.

In sum, we could show that it was possible to align different cell types with pre-defined distances by simply adjusting the G-Code. Especially, in Figure 2 e it is demonstrated that it is possible to print highly-dense cell-laden droplets with a partial overlap (220 μm group). To our knowledge this has not been done before with highly-dense cell-laden droplets and is not feasible for methods based on spheroids, which are already self-contained entities. Such partial cell contact may allow to directly compare behavior of cells in contact with a second cell type to cells in the same aggregate without contact to a different cell type. This is of particular interest to investigate developmental biological processes, such as vascularization, in which cells are constantly interacting with their microenvironment. The following sections describe results based on relevant cell types involved in such processes.

### 2.3. Influence of ASCs and HUVECs onto each other

After establishing and characterizing the process, red labeled ASCs and green labeled HUVECs were printed. They were either suspended in two separate cell suspensions or into the same bioink as a mixed cell suspension.

#### 2.3.1. ASCs and HUVECs as separate cell-laden droplets

N = 2 technical replicates with a total of n_full overlap_ = 11 cell aggregate pairs were printed and imaged daily for four days **(Figure 3 a)**. The distance between cell aggregate pairs were 4000 μm to avoid that they influence each other. On day two, HUVECs started to develop sprouts, whereas ASCs developed structures, which appeared similar to sprouts, and which were thus termed sprout-like structures. For each cell type, these structures grew larger in the subsequent two days. It is visible that ASCs did not inhibit sprout formation of HUVECs. This stands in contrast to recent findings, where the presence of spheroids containing human mesenchymal stem cells inhibited sprouting from HUVEC spheroids. ^[21]^ Differences may be explained by different cell sources for the stem cells and in the mentioned study the spheroids were not in direct contact with each other. Here, on the contrary, printing with a full overlap led to joint sprout-like structures on each sample, which appeared to be formed by HUVECs and ASCs together (indicated by the white arrows **Figure 3 b**). Such joint structures were detected on all cell aggregate pairs. It may be possible that HUVECs recruited ASCs, since it has been shown that ASCs can support the formation of vascular-like structures *in vitro*. ^[23]^ Nonetheless, most structures appeared to consist entirely of either HUVECs or ASCs instead of joint structures. Future studies could investigate this phenomenon and spatial arrangement of the cell types further.

**Figure 3:**
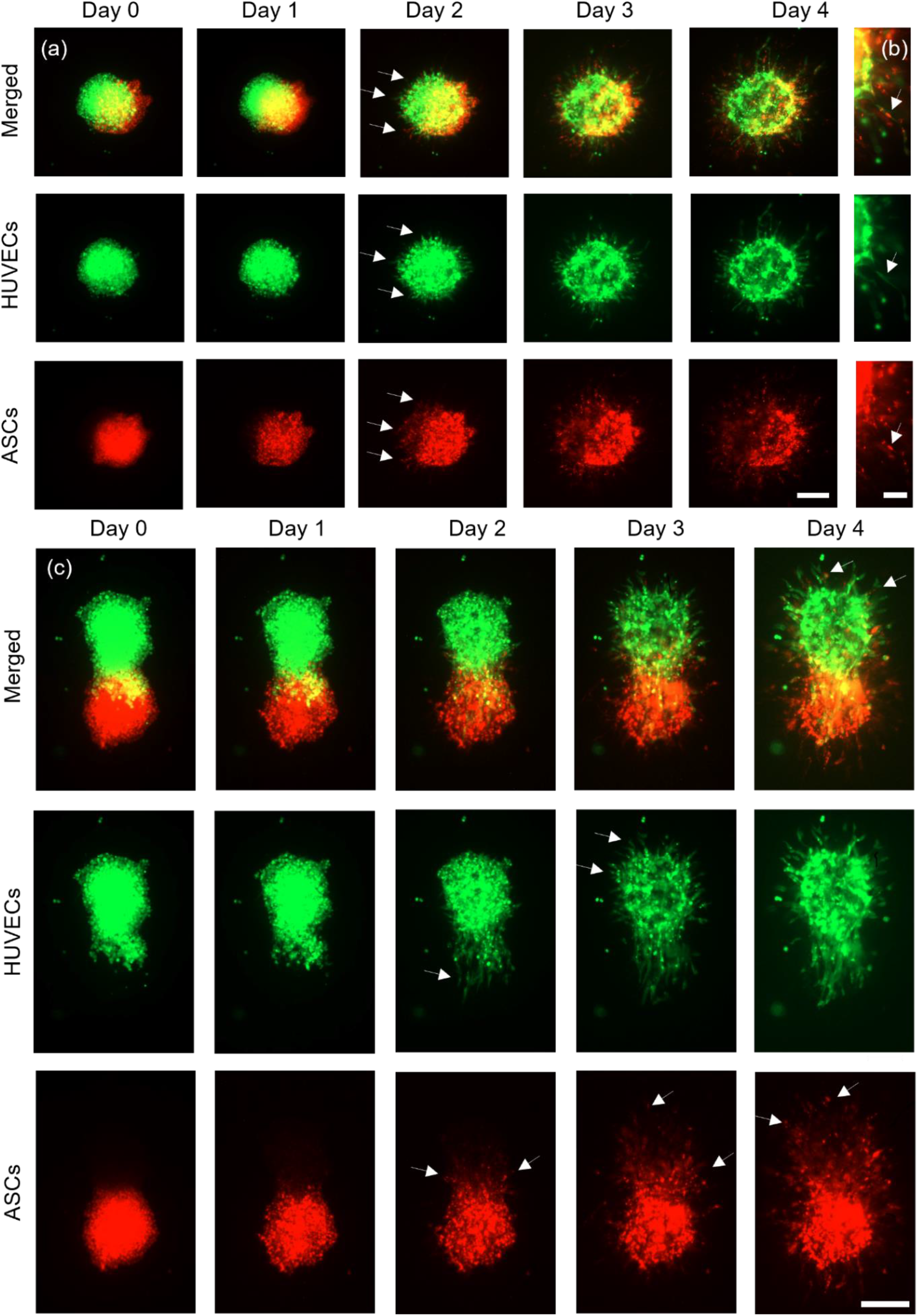
Representative images of ASCs and HUVECs. Aggregates printed with a full overlap (a, scalebar 200 *µm*) were imaged for four days and developed joint structures containing both cell types in the same structure (b, scalebar 100 *µm*). ASCs and HUVECs printed with a partial overlap are shown in (c) (scalebar 200 *µm*).

Next, ASCs and HUVECs were printed with a partial overlap (**Figure 3 c**) and imaged for four days. *N* = 2 constructs with a total of n_partial overlap_ = 11 cell pairs have been fabricated. On day two, the HUVEC aggregates formed clear sprout structures that appeared to permeate the ASC aggregates, as indicated by the white arrows Figure 3 c on day two. Interestingly, most of the HUVEC aggregates only showed clear sprouts in a region without ASCs one day later at day three (indicated by white arrows on Figure 3 c day three), which suggests that the direct cell-cell contact to ASCs may enhance sprout formation towards ASCs or that fibrin remodeling is more manageable around the ASC cell aggregate. ASCs also formed sprout-like structures and they appeared to proliferate preferably towards the HUVECs, as again indicated by the white arrows. On day four, ASCs could be found inside the HUVEC spots, where no signal was detected on day zero.

Last, ASCs and HUVECs were printed in proximity in groups with three different distances (distance between boundaries of the cell spots: 203 ± 24 μm, 372 ± 47 μm and 527 ± 57 μm).

For each distance, *N* = 2 technical replicates have been printed with a total amount of *n*_200 μm, ASCs_ = 6, *n*_200 μm, HUVECs_ = 6, *n*_370 μm, ASCs_ = 5, *n*_370 μm, HUVECs_ = 6, *n*_530μm, ASCs_ = 8, *n*_530 μm, HUVECs_ = 8 cell aggregates. Representative images for each group are shown in **Figure 4 a - c**. As described in the methods in detail, a region of interest (ROI) of 60° was defined towards the neighboring cell aggregates, and the CSL was measured in the ROI and the other region (300°). Afterwards, the results were normalized to 60° to achieve a better comparability. Measurements were performed on day three post-printing and results are shown in **Figure 4 d**.

**Figure 4:**
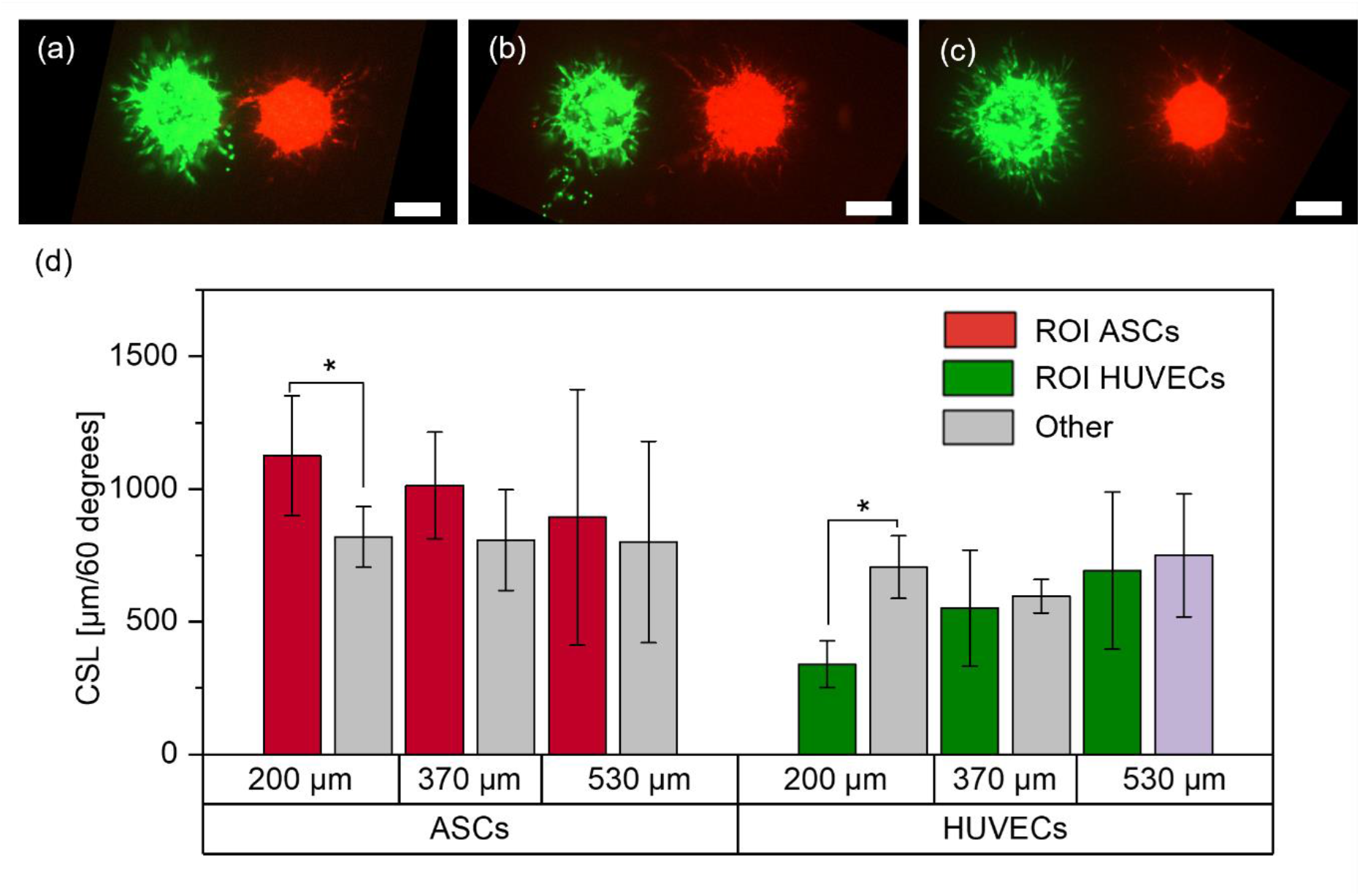
Representative images of ASCs and HUVECs in the 200 μm (a), 370 μm (b) and 530 μm (c) group on day three and respective measurements of the CSLs on day three (d).

For the 200 μm group, there is significantly less sprouting from HUVECs towards the ASCs, whereas there is significantly more sprout-like formation from the ASCs towards the HUVECs. This behavior decreases for increasing distances between the cell types and no significances were observed in the other groups.

Increased sprout-like formation of ASCs towards HUVECs is consistent with the results from the partial overlap, where ASCs grew and proliferated increasingly towards HUVECs. Released stimulation proteins (e.g. growth-factors) from HUVECs may recruit ASCs, which could also be one reason why the increased growth from ASCs towards HUVECs weakens for increasing distances. ^[24]^ Interestingly, while the behavior of ASCs is consistent for a gap between the cell types and a partial overlap, the behavior of HUVECs appears to change as soon as no cell contact between ASCs and HUVECs is established anymore. While there was clear sprout formation from HUVECs when printed with an overlap with ASCs, the opposite behavior was visible for close distances. These differences suggest that direct cell-cell contact may yield different behaviors of the cells in the aggregates. It is known that direct contact between the cells can change their behavior. For example, it has been reported that direct cell-cell contact between ASCs and HUVECs leads to an increased expression of αSMA or release of activin A. ^[24,25]^ Also, printing two cell types onto the same spot may lead to local differences in cell-hydrogel interactions which in turn may change cell behavior. Since the exact reasons for this behavior could not be identified, further research is required to understand the underlying causes. However, these findings can also be taken into account when designing larger tissue structures containing the applied cell types.

#### 2.3.2. ASCs and HUVECs as mixed cell suspension

After analyzing the behavior of ASCs and HUVECs printed as separate aggregates, their behavior was investigated, when mixed into the same bioink and dispensed as mixed cell-laden droplets. In analogy to a recent study, they were mixed with a ratio of 2 (HUVECs) to 1 (ASCs). ^[26]^ Mixed cell-laden droplets were printed with pitches of 500 μm, 800 μm, 1000 μm, and 2000 μm and imaged every day, for three days. Representative images of the cell aggregates for each distance and day are shown in **Figure 5 a – d**. Per pitch N = 2 technical replicates have been printed, except for the 1000 μm group with N = 3. In total, the number of corridors between cell-spots were *n*_500 μm_ = 42, *n*_800 μm_ = 26, *n*_1000 μm_ = 32 and *n*_2000 μm_ = 9. As a positive control for the behavior of HUVECs, HUVECs have been printed without ASCs with the same pitches. In total, N = 8 technical replicates with n_HUVECs_ = 54 aggregates have been printed. During the three days, mainly the ASCs developed sprout-like structures. In the 500, 800 and 1000 μm group, a clear directionality towards neighboring cell aggregates was visible. In the 500 μm group, already after one day, connections were formed with the neighboring cell aggregates. This trend was also clearly visible in the 800 μm and 1000 μm groups, where direct connections formed on day two and day three. A representative image of the 800 μm group on day three is shown **in Figure 5 e**. The influence of the pitch is further demonstrated by analyzing the CSL. As shown in **Figure 5 f**, the proportion of structures in the ROI decreases for increasing distances. For the 500 μm pitch, the CSL is more than 700 % higher in the ROI than in the other region on day one. The proportions decrease to about 300 % in the 800 μm group and about 60 % in the 1000 μm group on day one. Over the next two days these proportions slightly increased **(Figure S3)**. In addition, in the 2000 μm group, the CSL in the ROI was significantly higher than in the other region on days two and three whereas no significance was observed on day one. The structures formed by ASCs may be cellular bridges. Such bridges could form between ASC and ASC+HUVEC spheroids, as was recently reported. ^[18]^ These results also further support our previous findings that highly-dense cell-laden droplets may behave partially like spheroids. While the same experimental setup containing ASCs without HUVECs also showed directional growth of ASCs, no such bridges and no relation to the pitch were observed (**Figure S4**). The first stands in contrast to findings in which also pure ASC spheroids formed such bridges. ^[18]^

**Figure 5:**
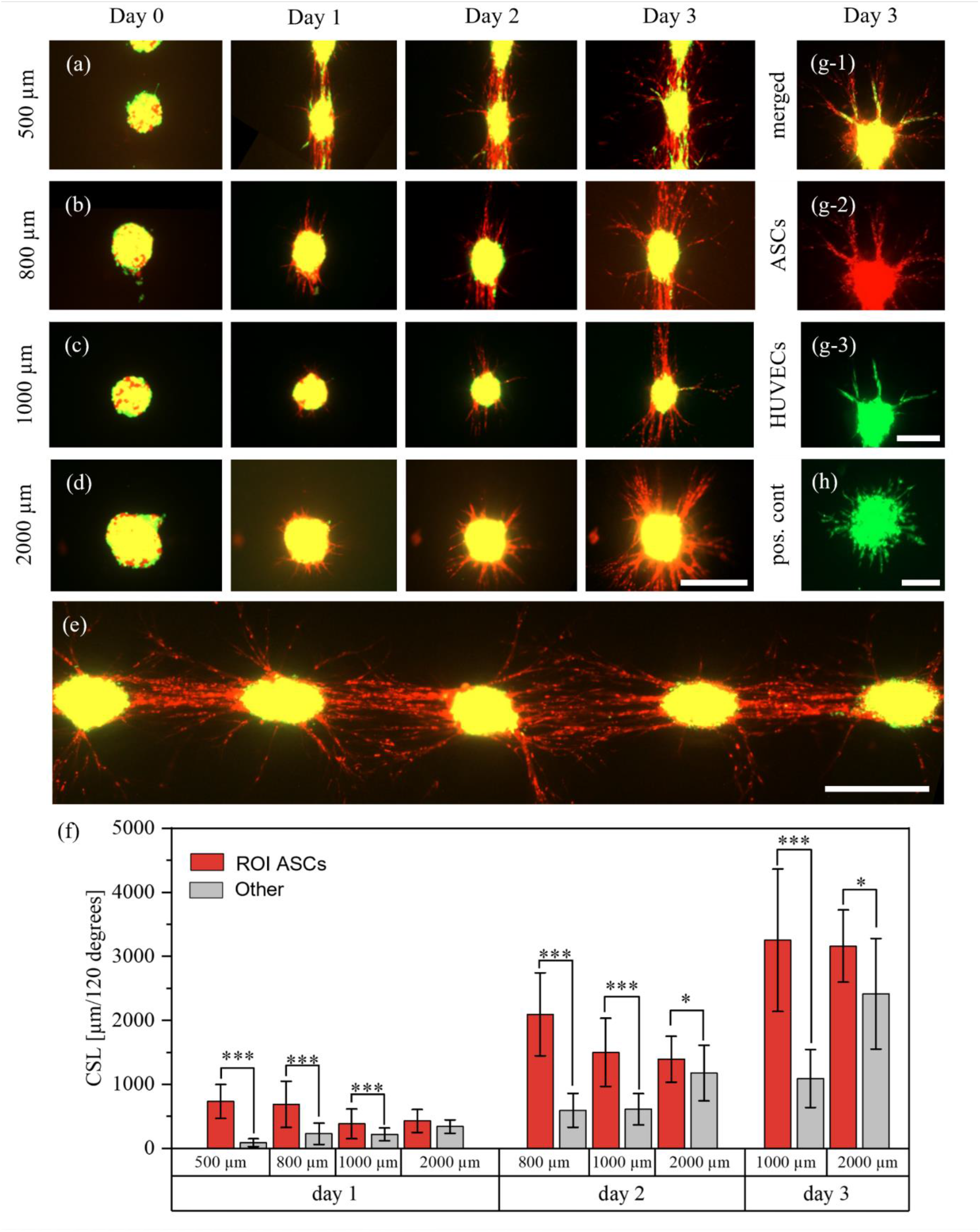
Behavior of ASCs and HUVECs in mixed aggregates. The cells were printed with different pitches and imaged for three days in which the ASCs formed cellular bridges between the aggregates in the 500, 800 and 1000 μm groups. A representative image of several cell spots on day three for a pitch of 800 μm is shown in (e). The structures of the ASCs were measured every day and are displayed in (f). However, on day 2 in the 500 μm group and on day 3 in the 500 and 800 μm group, no measurements were possible anymore. This was caused by too many cell structures which could not be separated visually anymore. A representative image of an aggregate where HUVEC sprouts were aligned along the ASCs is shown in (g). (h) displays the behavior of HUVECs in the positive control. Scalebars a-e: 400 μm. Scale bars g-h: 200 μm.

After the ASCs started to form sprout-like structures, HUVECs also occasionally developed sprouts. A representative image is shown in **Figure 5 g**. It is clearly visible that HUVEC sprouts aligned and organized themselves in dependence of the ASC structures. While 98 % of HUVEC aggregates in the positive control (pos. cont.) (**Figure 5 h**) developed homogenous sprouts with no preferred sprouting direction, only 45 % of HUVECs in mixed cell-laden aggregates developed sprouts. This difference in behavior may be caused by inhibitory effects of the stem cells onto HUVECs. ^[27]^ However, these observations stand in contrast to behavior of HUVECs and ASCs printed as separate droplets on top of each other as discussed above. These different behaviors may be caused by different interactions between the cells due to different cell distributions when mixing or printing on top of each other, but further research is required. As can be seen, ASCs formed more sprout-like structures than HUVECs, which is interesting since there were approximately twice as many HUVECs as ASCs in the bioink. However, since the applied method does not allow to analyze the spatial arrangement of the cells in three dimensions, exact statements can not be given. Also, the reason for formation of cellular bridges is not yet clear and further research is required.

### 2.4. Influence of SMC-differentiated ASCs and HUVECs onto each other

After the behavior of ASCs and HUVECs was analyzed, ASCs were substituted with ASCs that were differentiated into SMCs. This cell type was chosen due to their frequent use with endothelial cells for fabrication of vascularized structures as discussed above. The same experimental set-ups as with HUVECs and ASCs were chosen to evaluate similarities and differences between the behavior of the different cell types.

#### 2.4.1. Differentiation of ASCs

ASCs were differentiated to SMCs by TGF-beta1 treatment for 1 week. In relation to ASCs without TGF-beta1 treatment (**Figure 6 a**), TGF-beta1 induced the expression of alpha smooth muscle actin (αSMA), which is indicative for the differentiation of ASCs towards SMCs (**Figure 6 b**).

**Figure 6:**
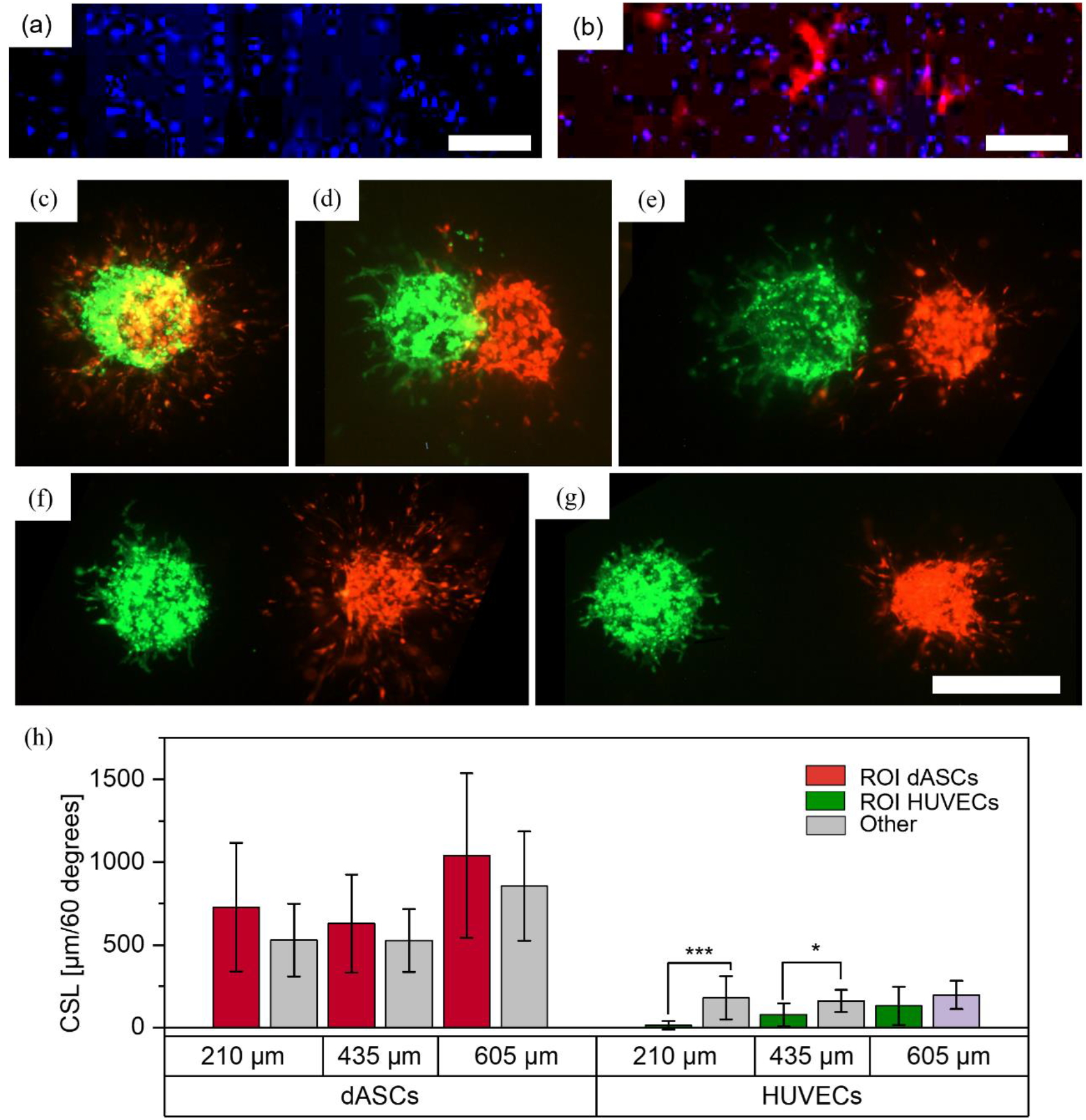
Printed patterns with dASCs and HUVECs as individual cell-laden droplets. ASCs were grown without TGF-β1 (a) and with TGF-β1 (b) and were stained with DAPI (blue) and with antibodies against αSMA (red).Scalebar: 500 μm. Aggregates were printed with a full overlap (c), partial overlap (d), and distances of approximately 210 (e), 435 (f) and 605 μm (g) between the boundaries of the aggregates. Scalebar: 500 μm. Measurements of the CSLs on day three are shown in (h).

#### 2.4.2. dASCs and HUVECs as separate cell-laden droplets

Differentiated ASCs (dASCs) and HUVECs were printed in similar experimental setups as undifferentiated ASCs and HUVECs (see chapter 2.3.1.). For a full overlap, N = 2 technical replicates with a total of *n*_*full overlap*_ = 8 cell pairs per distance were printed. A representative image on day three is shown in **Figure 6 c**. Again, as for ASCs and HUVECs, over the course of three days both cell types developed sprout, or sprout-like structures with no preferred direction for each cell type. For a partial overlap, N = 2 technical replicates with a total of *n*_*partial overlap*_ = 8 cell pairs were printed. A representative image is shown in **Figure 6 d**. While both cell types developed sprout(-like) structures again, in this constellation no growth of HUVECs towards the dASCs was observed which stands in contrast to findings from printing ASCs and HUVECs. This may be explained by different interactions of HUVECs in contact with SMCs and ASCs. Lastly, the two cell types have been printed with increasing distances and no contact in three different groups (distance between boundaries of the cell aggregates: 208 ± 26 μm, 435 ± 33 μm and 605 ± 30 μm, **Figure 6 (e – g**). N = 2 technical replicates with a total of *n*_*separate*_ = 8 − 15 cell pairs were printed and the CSLs have been analyzed on day three. For HUVECs, significant less sprouts formed in the ROI towards the dASCs in the 210 and 435 μm group. This behavior is consistent with observations from HUVECs and ASCs as discussed above.

In contrast to ASCs, HUVECs do not appear to influence the directionality of the sprout-like structures developed from the dASCs and no trend is visible. This was surprising as endothelial cells are known to recruit smooth muscle cells ^[28]^ Reasons for the anti directional formation of HUVEC sprouts are not known yet and require further research. However, as smooth muscle cells are often used with HUVECs for biofabrication of vascularized structures, this behavior can already be considered for possible bioprinting designs.

#### 2.4.3. dASCs and HUVECs as mixed cell suspension

dASCs and HUVECS were mixed (ratio 1:2) and printed with pitches of 500 μm, 800 μm, 1000 μm and 2000 μm. In total, N = 2 technical replicates (except 2000 μm group with N = 1) with n = 6 - 8 cell pairs per replicate have been fabricated. Images on day three are shown in **Figure 7 a-d**. As before, joint structures of both cell types were formed as indicated by the white arrows in **Figure 7 e**.

**Figure 7:**
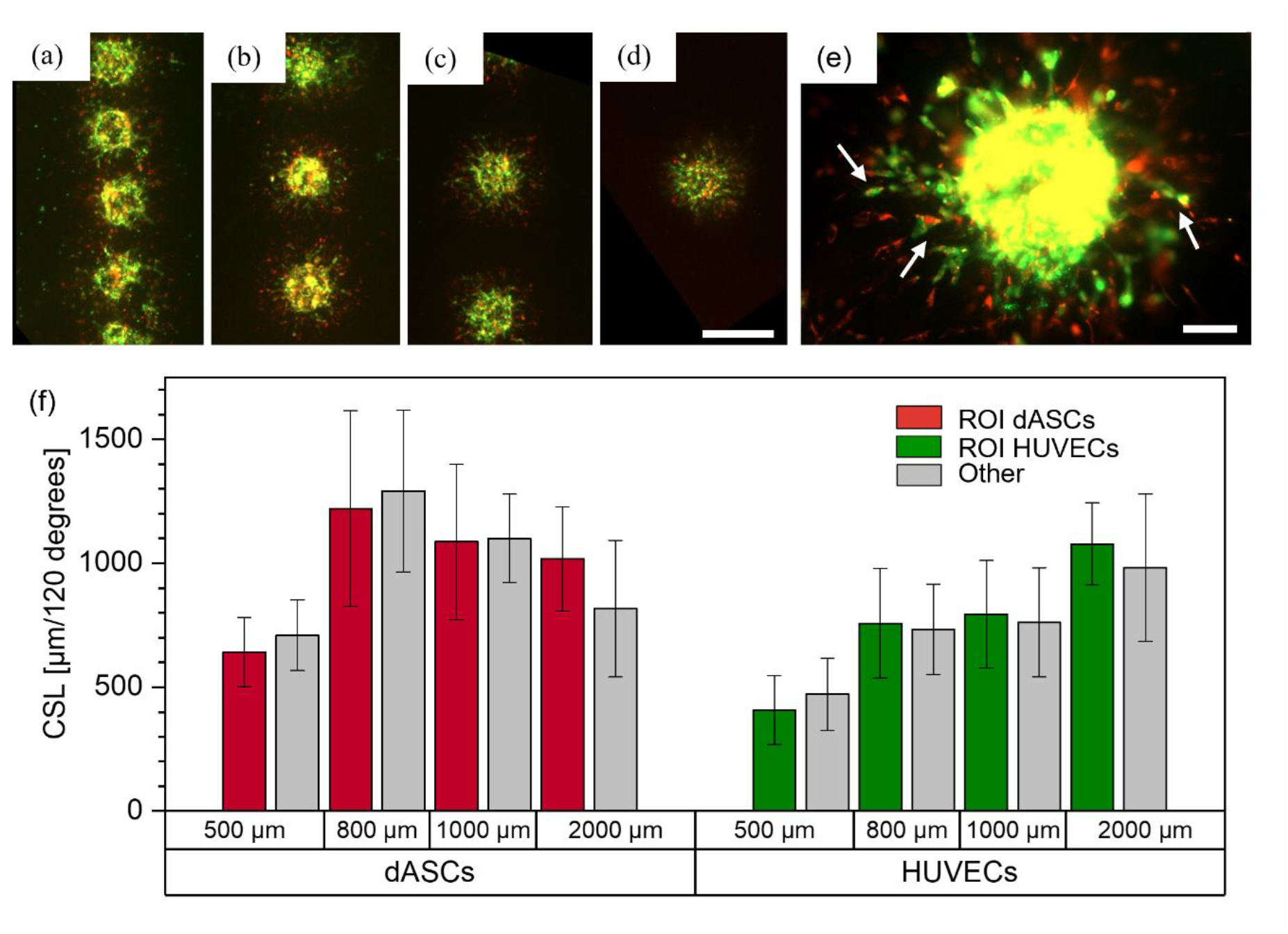
dASCs and HUVECs printed as mixed cell suspensions. Images for the 500 μm (a), 800 μm (b), 1000 μm (c) and 2000 μm (d) groups from day three are shown. Scalebar: 500 μm. (e) shows a mixed cell aggregate with joint sprout-like structures. Scalebar: 100 μm. (f) displays the measurements of the CSLs for each cell type and each group on day three.

In contrast to ASCs and HUVECs, no direct connections between the cell spots are visible, which may be caused by differentiation of ASCs into a different cell type. Also, all HUVECs displayed clear sprouting behavior, and not only few single sprouts as was the case for mixed cell suspension of HUVECs and ASCs. Thus, both cell types were analyzed for directed sprout formation towards the neighboring cell clusters and no directionality was observed (**Figure 7 f**). These observations suggest that in mixed cell suspensions each cell type has no inhibitory impact on the respective other cell type and that their post printing behavior does not necessarily depend on neighboring cell aggregates. Therefore, our findings indicate that bioprinting of pre-differentiated ASCs is beneficial for the creation of vascularized tissues. In the future, further methodologies such as confocal microscopy, immunostaining and gene expression analysis could be applied to analyze the spatial arrangement of the individual cell types.

## 3. Materials and Methods

### 3.1. Process to print different cell types with high precision on the same substrate

A bioprinter which is equipped with a drop-on-demand dispenser and a robotic stage that can move in x-, y-, and z-direction was used for all experiments (Figure S5). To prepare the bioink, each cell type was suspended individually into fibrinogen to obtain a concentration of approximately 200 – 250 cells per 10 nl droplet. Fibrin hydrogels were prepared on 18×18 mm glass slides and were crosslinked in a humid atmosphere at 37 °C. Bioink containing the first cell type was transferred into the dispenser and the volume of the droplet was calibrated to 10 nl via an integrated optical system (SmartDrop, Biofluidix GmbH) that is installed in the bioprinter. The dispenser with the first bioink was left for 15 minutes during which the concentration increased to about 700 cells per 10 nl due to sedimentation (Figure S6) while the other bioink was kept on ice. During the 15 minutes a droplet was ejected every three seconds to prevent clogging of the dispenser. After 15 minutes, glass slides with prepared hydrogel substrates were transferred onto a customized platform that contained notches which exactly fit the glass slides (Figure S5). Then the bioink was printed onto the fibrin samples in form of a predefined pattern that was set via G-code. If a second bioink with a second cell type was used, after printing the first cell type onto the samples, the samples were re-transferred into a humid atmosphere to prevent drying of the hydrogel and the second cell type was filled into a new capillary. After leaving the second dispenser for 15 minutes while ejecting a droplet every three seconds, the samples were transferred onto the notches of the platform again and the second bioink was printed in relative positions to the cell aggregates of cell type one. Afterwards the samples were incubated for 60 minutes during which the cell clusters adhered to the hydrogel due to enzymatic crosslinking of the fibrinogen in the bioink and the thrombin from the fibrin layer. Subsequently, a second layer of fibrinogen was added to create a sandwich-like structure and the constructs were incubated in endothelial cell growth medium (ECGM). During incubation in the following days, the cells developed structures which could be analyzed further, for example via microscope or antibody staining, to assess the influence of the different cell types onto each other, e.g. enhanced growth towards a neighboring cell aggregate.

### 3.2. Cell culture

All cell types were maintained at 37 °C with 5 % CO_2_ in a humidified atmosphere using the specific cell culture medium. Passaging of cells was carried out when cells reached a confluency of less than 90 %. After cells were harvested and samples were printed, all samples were incubated in ECGM. Cells were stained with different CellTrackers™ to distinguish between the cell types.

#### 3.2.1 HUVECs

Primary human umbilical vein endothelial cells (HUVECs) that were isolated from the vein of the umbilical cord of a single donor (PromoCell GmbH, C-12200, Germany) were cultured in endothelial cell growth medium (ECGM) (PromoCell GmbH, C-22010) with 1 % penicillin/streptomycin (P/S) (Life Technologies, 15140-122) and 10 % fetal calf serum (FCS) (10270-106). A supplement mix containing different factors was provided by PromoCell and added to the culture medium. HUVECs from passages three to six were used

#### 3.2.2. Differentiated and undifferentiated ASCs

Adipose tissue derived stem cells (ASCs) were isolated from a 24-year-old male donor using an established protocol, as described before.^[29]^ They were cultured in EBM™-2 basal medium (LONZA, CC-3156) that was supplemented with EGM™-2 SingleQuots™ supplements (LONZA, CC-4176), 10 % FCS and 1 % P/S. ASCs from passage two to seven were used. Also, ASCs were differentiated (dASCs) to smooth muscle cells (SMCs) according to a protocol that was reported previously.^[30]^ In brief, at 90 % confluency, ASCs were incubated in minimum Essential Medium α (α-MEM) (GibcoTM MEM α, Nukleoside, GlutaMAXTM Supplement, 32571028) (with 1 % P/S and 10 % FCS) which was supplemented with TGF-β1 (c_TGF_ = 5 ng/ml) for seven days. Cell culture medium was refreshed at day three and day six. DASCs from passage three to seven were used.

#### 3.2.3 Immortalized mesenchymal stem cells

Immortalized mesenchymal stem cells (iMSCs) ^[31]^ were cultured in α-MEM with 1 % P/S and 10 % FCS.

### 3.3. Bioink preparation

To fabricate the bioink, cells were harvested and pelleted at 1,000 rpm for 5 minutes. For HUVECs and iMSCs, fibrinogen (10 mg/ml) was mixed with ECGM 1:1 and the cells were suspended in the ink to obtain a concentration of 25*10^6^ cells/ml. For ASCs and dASCs, fibrinogen (10 mg/ml) was mixed with PBS 1:1 and the cells were suspended in the ink to obtain a concentration of 20*10^6^ cells/ml. The fibrinogen in the bioink increased the adhesion of the cell spots due to crosslinking of fibrinogen in the bioink and the thrombin present in the fibrin hydrogel layer.

### 3.4 Hydrogel Matrix

Fibrin was chosen to embed the cell aggregates in a 3D environment. It was used as a substrate layer as well as covering layer after printing, thus resembling a sandwich structure. It was prepared by mixing fibrinogen (10 mg/ml) and thrombin (10 mg/ml) in a 1:1 ratio. Fibrin was prepared on 18×18 mm glass coverslips and samples were incubated at 37°C in a humid atmosphere for 30 minutes before the printing process was started.

### 3.5 Drop-on-demand technology

After preparing the bioinks, they were transferred to a piezoelectric drop-on-demand dispenser (PipeJet Nanodispenser, Biofluidix GmbH, more details were described previously ^[32]^) with exchangeable capillaries with a 200 μm diameter. The dispenser was connected to a reservoir in which the bioink (between 50 and 150 μl) was filled by capillary forces. The dispenser was connected to a bioprinter with a three-axis robotic stage which also had an integrated optical system (SmartDrop, Biofluidix GmbH) to measure the shape of the dispended droplets in-flight. Based on the shape of the droplet, the volume could be calculated and the process parameters adjusted accordingly to dispense a volume of 10 nl.

### 3.6 Printing design and quantification of Results

For this work two printing designs were used: printing aggregates of two cell suspensions next to each other or printing several aggregates from the same cell suspension in a line **(Figure 8 a)**. The position between the cell types was described by either the pitch p (distance between the aggregates’ centers) or distance d (distance between the boundaries of the aggregates). A ROI was defined, which was a window of 60° towards the other cell type for cell pairs or two windows of 60° towards the neighboring cells for a line pattern. Thereafter, the images were edited with imageJ and skeletonized (**Figure 8 d**). The applied method yields a projection of a 3D dimensional construct, thus only statements towards the imaged planes can be made. This was sufficient to determine the relative growth of cell types in the ROI and “other” areas. Usually at least two technical replicates with at least biological triplets were created for each experiment. However, depending on the print design, the number of aggregates varied between three and twenty cell spots per technical replicate. Hence, this information is given for each experiment individually. After skeletonizing, the CSL was measured in the ROI and the other region using imageJ. As the ROI was smaller than the other region, the results from the other region were normalized to the ROI. Afterwards, a paired student’s t-test was used to determine statistical significance between the ROI and other region. A probability value of *P < 0.05 indicated statistical significance, with increasing significances for **P < 0.01 and ***P < 0.001.

**Figure 8:**
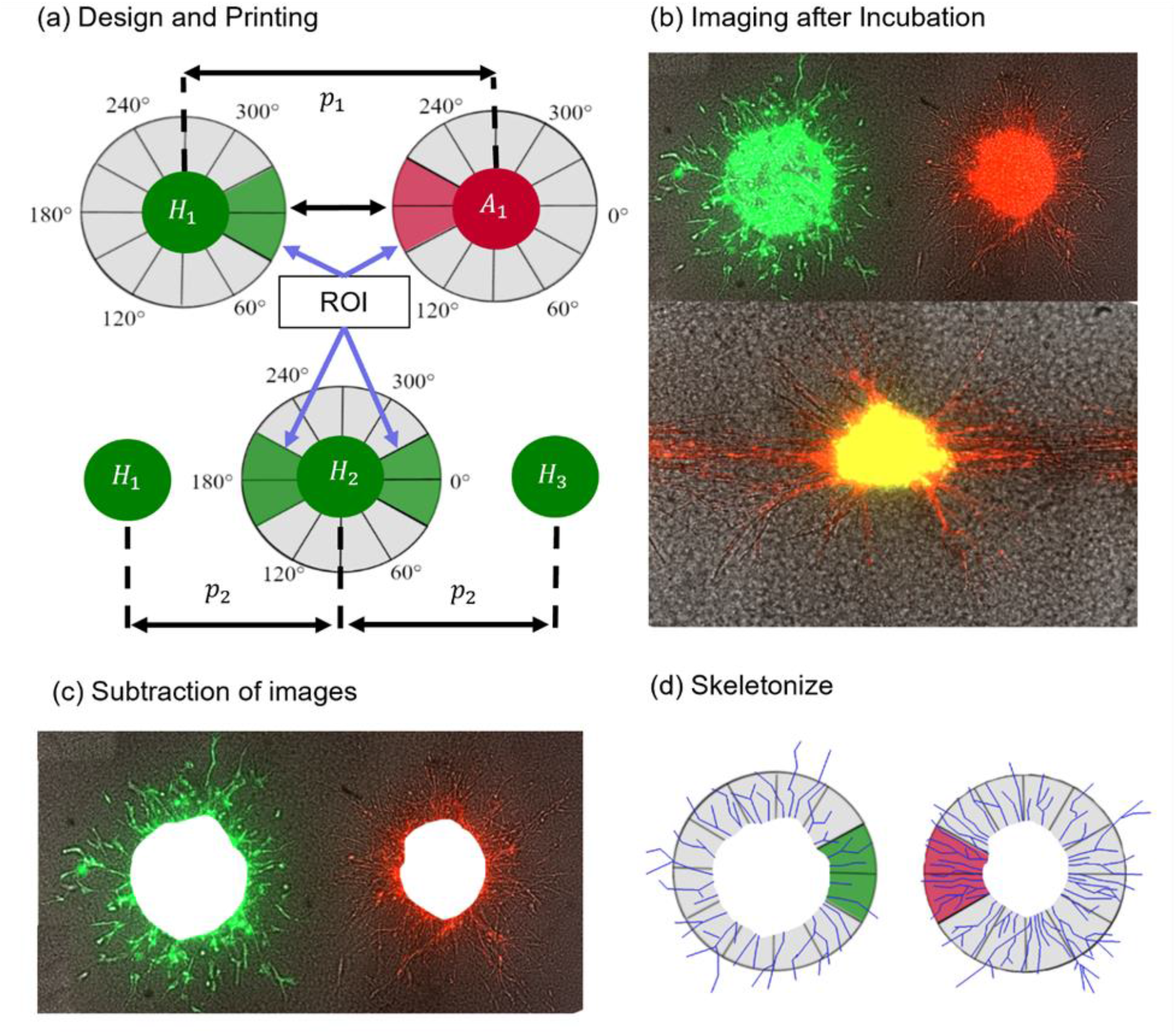
Schematic of the steps from printing design to analysis. First one of the print designs is fabricated (a), then aggregates are imaged after several days of incubation (b). Next, the images from day 0 were substracted (c) and the aggregates were skeletonized to analyze the structures in the ROI and other region.

### 3.7 Labeling and imaging of cells

CellTracker™ was used to stain cell types differentially. CellTracker™ RED CMTPX dye (ThermoFischer Scientific, C34552) was used to label either ASCs or differentiated ASCs, whereas CellTracker™ Green CMFDA Dye (ThermoFischer Scientific, C2925) was used to label HUVECs. Both dyes were dissolved in dimethyl sulfoxide (DMSO) (CAS 67-68-5, Sigma-Aldrich), and a stock solution with a final concentration of 10 mM was obtained and stored at -20 °C. The working solution was prepared by diluting the stock solution to a final concentration of 5 μM in cell-specific serum-free cell culture medium. After cells were harvested and pelleted, the pellet was suspended in 2 ml of the working solution and incubated for 40 min in 37°C. Afterwards, the cells were washed, centrifuged, the supernatant was removed and the cells were suspended into the bioink.

### 3.8 Live-dead assay

To assess influence of the printing process onto the viability of the cells, a live-dead assay was performed immediately after encapsulating the cells with the second fibrin layer. Samples were stained by a PBS-based staining solution of 2 μM Calcein-AM from Thermo Fischer Scientific (Waltham, USA) for viable cells and 6 μM Ethidium homodimer-1 from Sigma Aldrich (St.Louis, MO, USA) for dead cells for 30 min at room temperature in the dark. Cells pipetted manually and not printed via DoD were used as the positive control. For the negative control, cells were immersed in 80 % ethanol for 30 min. Fluorescence images were obtained with excitation wavelengths of 470 nm for Calcein-AM and 555 nm for Ethidium homodimer-1. To ensure a consistent evaluation, an approach as previously published was used to assess the signals. ^[33]^ The signals were transformed into rainbow-pseudo colors and only signals with intensity values higher than 70 % of the maximum intensity were taken for further analysis. Finally, the cell viability was calculated by dividing the number of viable cells through the number of total counted cells.

## 4. Conclusions

To fully yield the potential of future bioprinting approaches there is an increasing demand to observe how cells influence each other in 3D environments and to understand which role the distance between the cell types may play. However, only few approaches allow to precisely position highly-dense and locally confined cell aggregates in 3D. Here, a novel approach based on a sandwich-technique was introduced, which allows deposition of highly-dense cell-laden droplets from two different cell types with high precision on a fibrin hydrogel. Average alignment errors between runs were approximately 250 μm and errors within runs were less than 60 μm. ASCs were printed with HUVECs as individual cell aggregates and significant less sprouting from HUVECs towards ASCs was observed for a gap of 200 μm between them, whereas ASCs formed significantly more structures towards HUVECs for that distance. Partial reduced sprouting from HUVECs towards ASCs is particularly interesting because it has been reported that the presence of a MSC (of which ASCs are a subgroup) spheroid in close proximity to a HUVEC spheroid leads to an entire inhibition of HUVEC sprouts. ^[21]^ Directed growth of ASC structures towards HUVECs may be explained by recruitment of ASCs by HUVECs, which has been demonstrated previously. ^[24]^ For mixed cell suspensions, ASCs formed directed structures and cellular bridges between the aggregates for pitches up to 1000 μm and potential inhibitory effects of ASCs on HUVECs were observed. Such bridges were also recently reported for ASC+HUVEC spheroids for inter spheroid-distances up of 200 μm. ^[18]^ These distance differences and the mechanisms leading to bridge formation require further investigation. The same experiments were repeated with ASCs that were differentiated in SMCs. For these cell types, significantly less sprouts from HUVEC aggregates towards dASCs was observed, whereas the HUVECs appeared to have no measurable impact on dASCs. For mixed cell-suspensions, no directionality of any cell type was observed and all HUVECs displayed good sprouting behavior. For ASCs, as well as dASCs, combined with HUVECs, often joint structures containing two cell types was observed.

These findings demonstrate the capabilities of the developed method to discover new phenomena for the interaction between different cell types. In the future the developed approach should be combined with other methodologies to explain the causes for the observed cell behaviors, which are not fully understood yet. Nonetheless, directional and anti-directional growth of cell types could already be considered when biofabricating tissues and structures with the investigated cell types and the method could be also applied to investigate the interactions between other cell types.

## Data Availability Statement

All data needed to evaluate the conclusions in the paper are present in the paper and/or the Supplementary Materials. Additional data related to this paper may be requested from the authors.

## Acknowledgements

The authors thank B. Baumer and L. Rigger for excellent technical assistance. Prof. Dr. Matthias Schieker from the Laboratory of Experimental Surgery and Regenerative Medicine of the Ludwig Maximilians-University of Munich for providing the immortalized cell line. The microscope icon in figure 1 has been designed using resources from flaticon.com. This work was supported by funding of the Deutsche Forschungsgemeinschaft (FI790/10-2, and KO3910/1-2).

## CRediT authorship contribution statement

Joshua Weygant: Methodology, Investigation, Visualization, Writing – original draft, conceptualization. Fritz Koch: Validation, Writing – review and editing, supervision. Katrin Adam: Investigation. Kevin Troendle: Validation, Writing – review and editing. Roland Zengerle: Supervision, conceptualization. Guenter Finkenzeller: Resources, writing – review and editing. Sabrina Kartmann: Formal analysis, Writing – review and editing, resources, Peter Koltay: Formal analysis, Conceptualization, Validation, Writing - review & editing. Stefan Zimmermann: Formal analysis, Conceptualization, Validation, Writing - review & editing, project administration, Funding acquisition

## Declaration of competing interest

The authors declare that they have no known competing financial interests or personal relationships that could have appeared to influence the work reported in this paper.

## Notes

### Competing Interest Statement

The authors have declared no competing interest.

