## Supplementary material for "A drop-on-demand bioprinting approach to spatially arrange multiple cell types and monitor their cell-cell interactions towards vascularization based on endothelial cells and mesenchymal stem cells": SI_Information

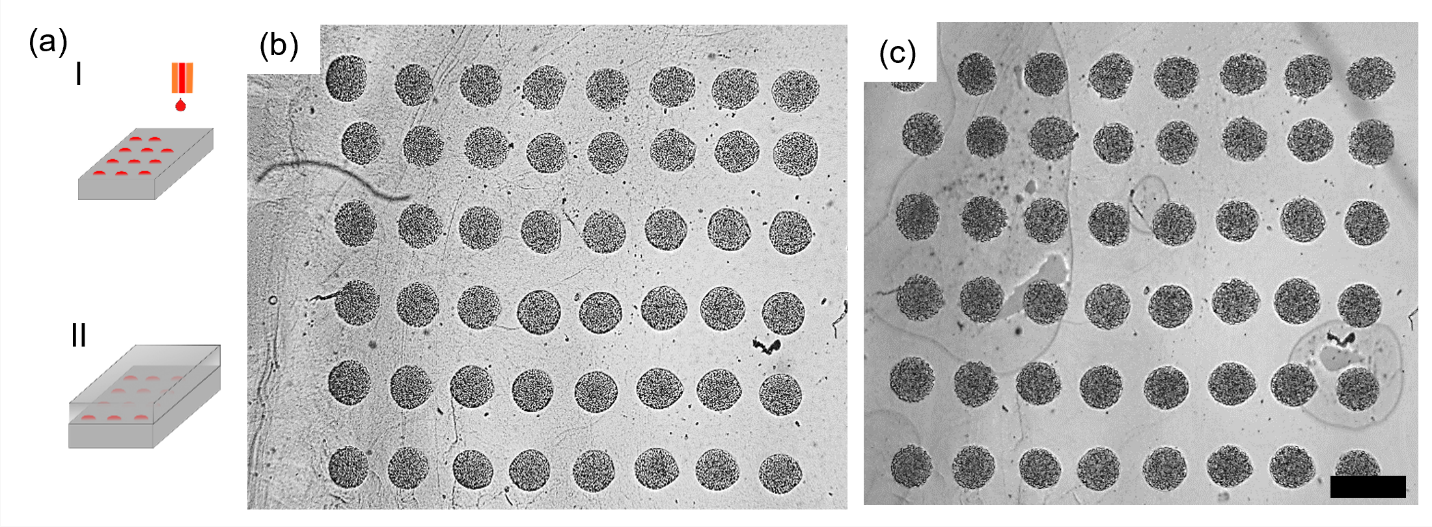

Figure S1: Experiment to determine shape fidelity by bioprinting of iMSCs arrays in 3D environments. (a): Schematic of the experimental setup. IMSCs were printed onto a fibrin substrate, another fibrin layer was added, and the substrate was analyzed optically. (b): Printed patterns of iMSCs on the first fibrin layer before adding the second fibrin layer. (c): Printed pattern after adding the second layer of fibrin. Scalebar: 500 μm.

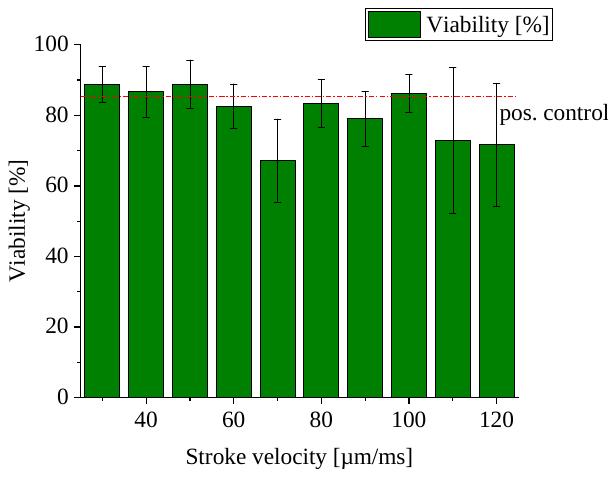

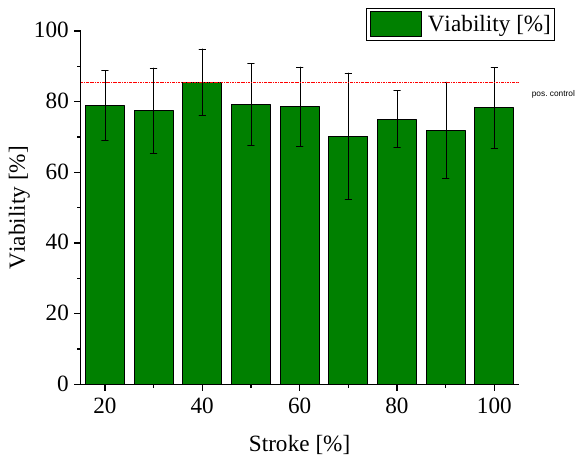

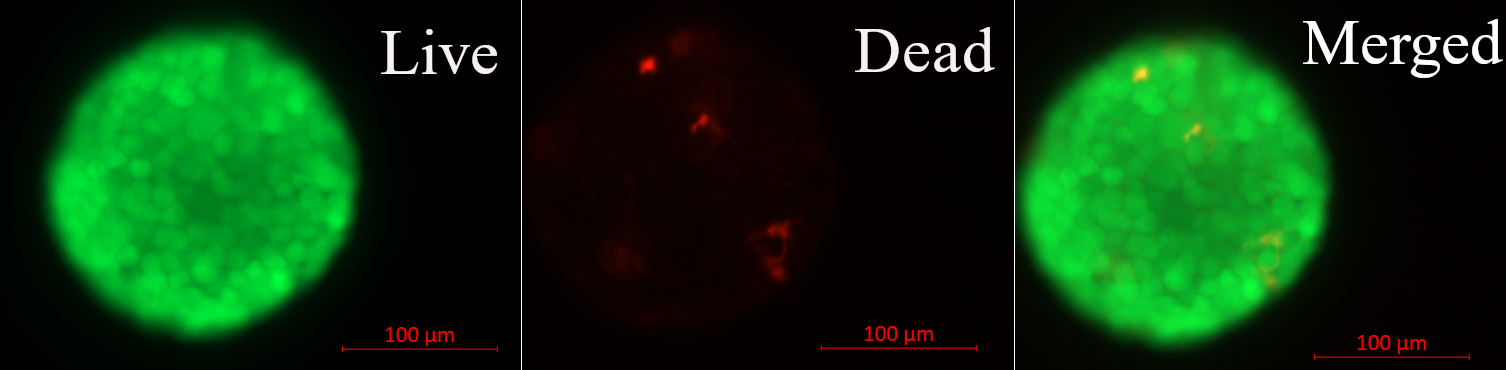

Live

Dead

Merged

(a)

(b)

(c)

Figure S2: (a) Representative fluorescence images from a cell aggregate after printing and staining with live/dead assay. Separate images, as well as merged image, are shown. (b): Measurement diagram showing the viability of iMSCs post-printing for different strokes and constant stroke velocity. (c) measurement diagram showing the viability for different stroke velocities with constant stroke

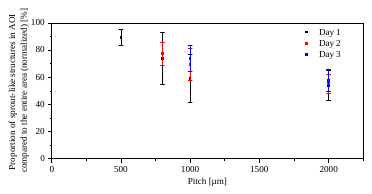

Figure S3: Proportion of ASC structures in the ROI compared to the entire area over the course of three days. On day 2 in the 500 μm group and on day 3 in the 500 and 800 μm group, no measurements were possible anymore. This was caused by too many cell structures which could not be separated visually anymore.

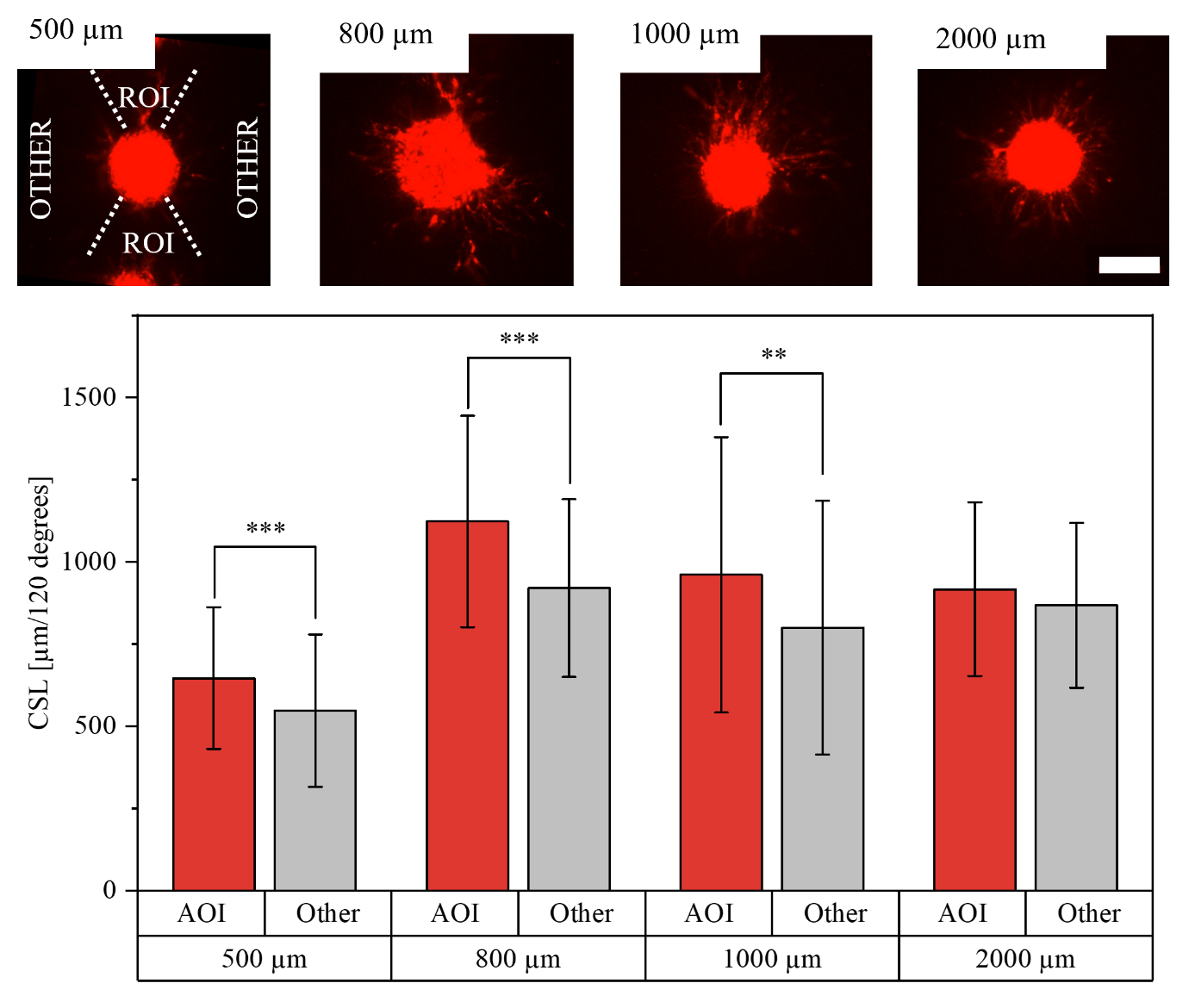

Figure S4: Top: Fluorescence images of ASC-laden droplets with different pitches from their neighbors, which are above and beneath the aggregates, as indicated on the 500 μm image by the "region of interest (ROI)" and the region "other" with no cell aggregates at day four in fibrin. Scale bar: 200 μm. Bottom: Measurement of mean values and standard deviations of cumulative sprout length (CSL) in the ROI compared to the region with no cell aggregates on day four. (*P < 0.05, **P < 0.01, ***P < 0.001). N = 4 technical replicates have been printed and in total n_500_ = 64, n_800_ = 42, n_1000_ = 28, n_2000_ = 16 corridors between ASC aggregates have been analyzed.

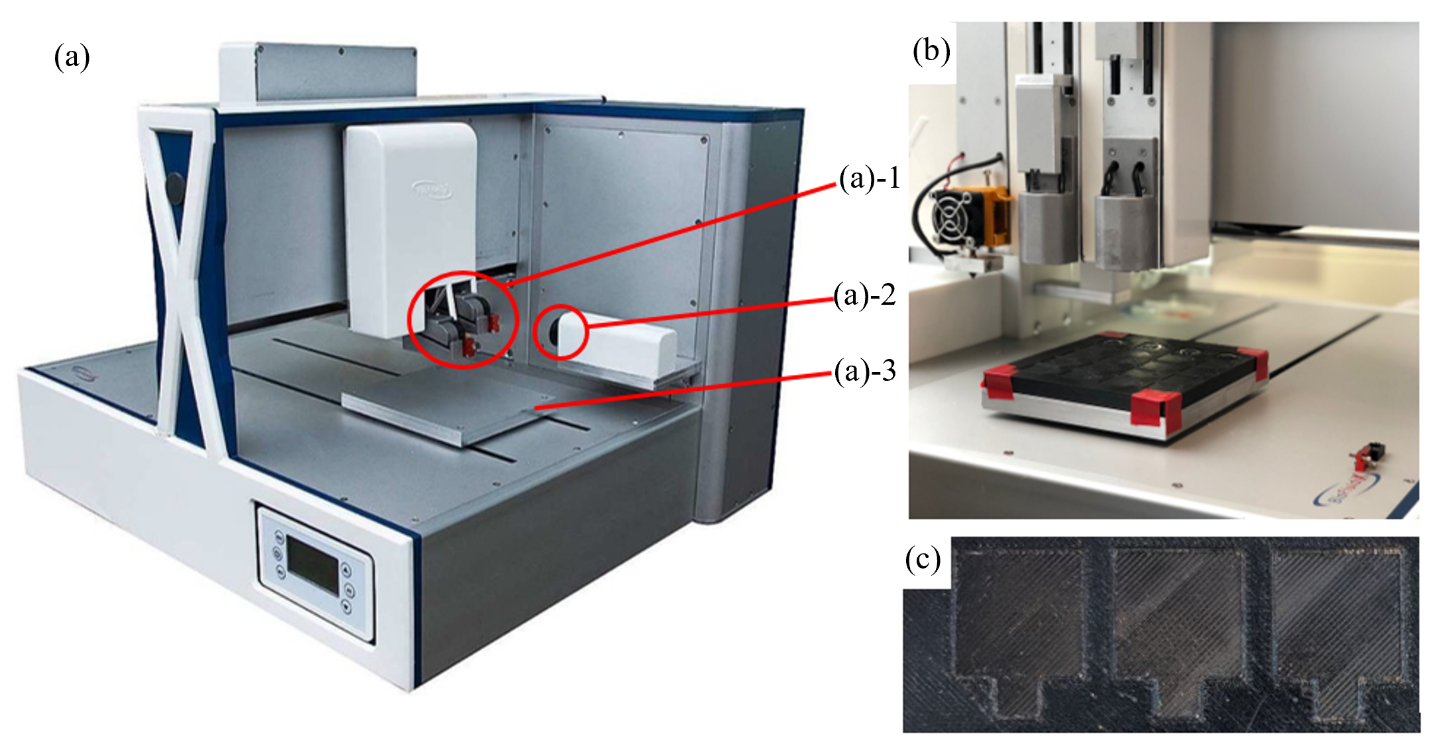

Figure S5: Used Bioprinter and printing platform. The bioprinter (a) is equipped with two drop-on-demand dispensers (a-1), a camera to capture the ejected droplets (a-2) for the SmartDrop system and a movable platform (a-3). A customized platform was 3D printed and attached to the platform (b) of the printer. The customized platform had notches (c) which exactly fit to the used glass slides to precisely place the substrates into the same position repeatedly.

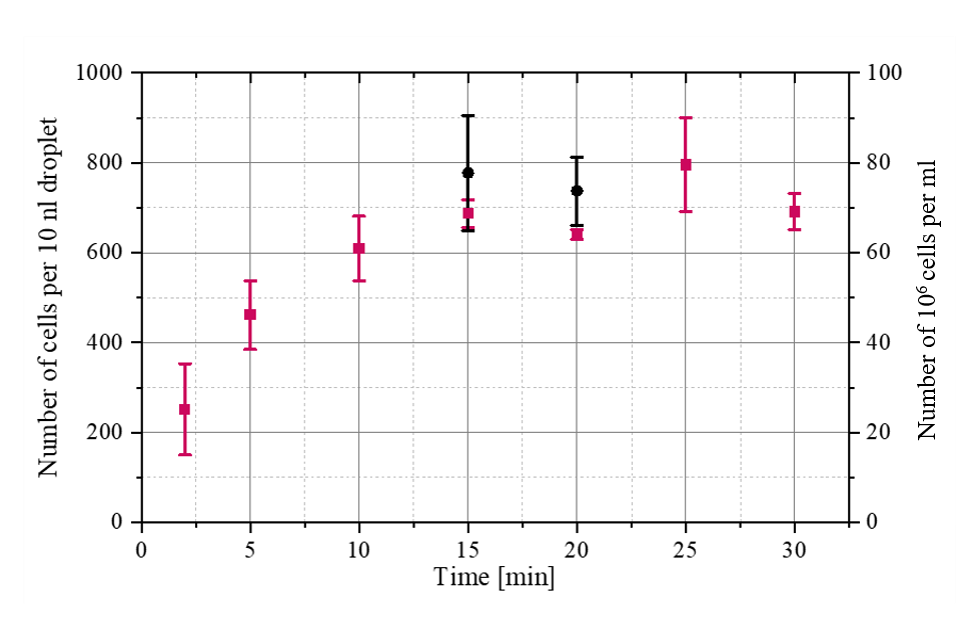

Figure S6: Sedimentation of ASCs for two experiments. First, ASCs were loaded into fibrinogen with a concentration of approx. 200 cells/10nl and a droplet was ejected every three seconds to prevent clogging of the dispenser. Every five minutes, first 10 cell-laden droplets were dispensed and discarded and then 100 cell-laden droplets with a volume of 10 nl were dispensed into two Eppendorf tubes (with 50 droplets per tube) and cells were counted with an automatic cell counter (Countess II, ThermoFischer Scientific). Per tube, two measurements have been performed. Results are shown in form of the red measurement points above. This experiment was repeated, but 100 cell-laden droplets were only ejected at 15 and 20 minutes. Results are shown in form of the black measurement points above. The latter experiment was repeated for HUVECs and concentrations of approximately 700 cells/10 nl droplet after 15-20 minutes were measured.
